# The role of the Frontal Aslant tract in bilingual language control

**DOI:** 10.1101/2023.02.01.526563

**Authors:** Cinzia Quartarone, Eduardo Navarrete, Simone Gastaldon, Sanja Budisavljević, Francesca Peressotti

## Abstract

In the present study, we investigated the microstructural properties of the right and left Frontal Aslant tract (FAT) in relation to bilingualism and language modality by comparing a group of unimodal bilinguals (i.e., bilinguals proficient in two spoken languages) and a group of bimodal bilinguals (i.e., bilinguals proficient in a spoken and a signed language). We found that the microstructural properties of the left FAT were related to the performance in semantic fluency in the second language (L2), either signed or spoken. Additionally, only for bimodal bilinguals, the microstructural properties of the right FAT were related to picture naming performance in the first spoken language (L1). No significant effects on performance were found in a language comprehension task. Overall, the results suggest that the FAT plays a significant role in language production in bilinguals. The left FAT appears to be involved primarily during the use of spoken or signed L2, while the right FAT appears to be involved in handling the competition of the signed L2 language while speaking L1.

## 1. INTRODUCTION

The frontal aslant tract (FAT) is a recently described intralobar frontal tract (Catani et al., 2012; Thiebault de Schotten et al., 2012) connecting the supplementary motor complex in the superior frontal gyrus (comprising the supplementary motor area - SMA - and the pre-SMA) to the most posterior part of Broca’s area, the lateral IFG (see Figure 1). Based on established functions of connected regions, it has been hypothesized that the FAT plays a role in control and executive functions, in particular in inhibitory control during speech production (Shekari & Nozari, 2023; Ribeiro et al., 2024). In the present study, we focus on two bilingual populations, bimodal bilinguals (L1 spoken, L2 signed) and unimodal bilinguals (L1 spoken, L2 spoken), examining whether their linguistic performance is associated with the microstructural properties of the Frontal Aslant tract (FAT). Importantly, this is the first study investigating the FAT in relation to bilingualism.

**Figure 1.**
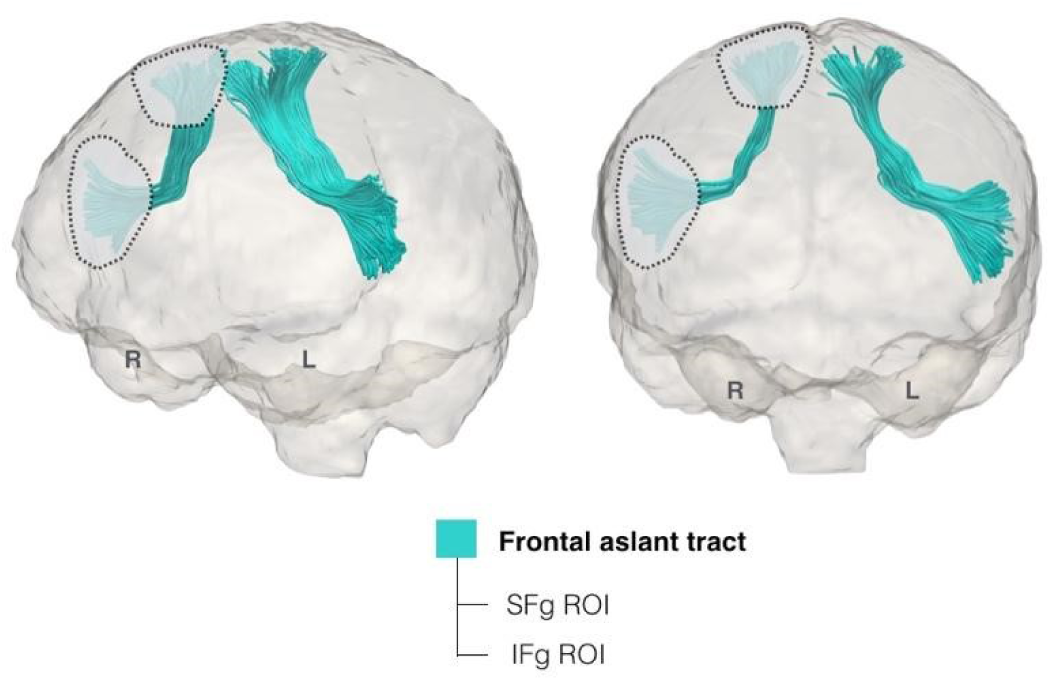
Tractography reconstructions of the left and right FAT in a representative subject. Frontal ‘AND’ ROIs including superior frontal gyrus (SFg ROI) and inferior frontal gyrus (IFg ROI) are shown for the right hemisphere.

In a recent review, Dick et al. (2019) proposed that the FAT is a key component of a neural circuit that, together with subcortical and cerebellar regions, is engaged in action control, specifically in planning, timing, and coordination of sequential motor movements. Within this network, the FAT would not be simply involved in motor processing but would perform a domain-general higher-level function, i.e., resolving the conflict among competitor motor programs. In the same review, Dick and coll. proposed some degree of hemispheric specialization. While the left FAT is considered part of a circuit specialized for speech action control (Tremblay & Dick, 2016), the right FAT is considered part of a circuit specialized for general action control. Neuroimaging studies have extensively evidenced the importance of the areas connected by the left FAT in language processing. The left IFG appears to be implicated in controlled semantic and lexical retrieval both in spoken and signed language (e.g. Katzev et al., 2013; Emmorey et al., 2007), and the SMA and pre-SMA are associated with selecting and executing motor programs for speech (Alario et al., 2006; Tremblay & Gracco, 2009; 2010), as well as for other non-linguistic domains (Cona & Semenza, 2017). Empirical evidence supporting the involvement of the left FAT in language production comes mainly from studies in various clinical populations, which report a relationship between speech fluency and the microstructural properties of the left FAT (e.g., Catani et al., 2012; Mandelli et al., 2014; Alyahya et al., 2020; Dragoy et al., 2020; Li, M. et al., 2017). Furthermore, intraoperative stimulation studies have shown that direct stimulation of the left FAT often results in speech arrests (see e.g., Fujii et al., 2015; Kinoshita et al., 2015). Other studies suggested an association between the FAT and stuttering: The severity of persistent developmental stuttering seems to correlate with the diffusivity of the FAT (Kronfeld-Duenias et al., 2016). When considering studies on non-clinical populations, evidence shows that the microstructural properties of the FAT are associated with language tasks not directly related to fluency. Broce et al. (2015) found a correlation between the length of the left FAT and the score on a receptive language battery in typically developing children 5-8 years of age. Vallesi & Babcock (2020) reported that the degree of left asymmetry of the FAT was correlated with lexical decision speed but not with verbal fluency performance in a group of healthy university students (see also Kronfeld-Duenias et al., 2016, who found no correlation between the properties of the FAT and the fluency task in non-clinical adults).

Regarding the right FAT, functional neuroimaging studies have shown activation of both the right SMA and pre-SMA and the right IFG in tasks requiring stopping behaviors (e.g., Nachev et al., 2008; Garavan et al., 1999). Stimulation of these areas has been associated with inhibiting voluntary fine movements (Luders et al.,1988). Patients with SMA and pre-SMA lesions exhibited deficits in producing complex sequences of movements (Dick et al., 1986), and patients with lesions in the IFG show impairments in inhibiting irrelevant task sets (Aron et al., 2003; 2004). However, patients with right FAT resection do not show impairments in the performance of the Stroop test, where inhibition of verbal responses is required (Puglisi et al., 2019). This further supports the hypothesis of at least partial hemispheric specialization of the control network involving the FAT.

This body of evidence aligns with the proposal that the FAT is involved in inhibitory control processes, particularly for those related to the regulation of speech output (Shekari & Nozari, 2023). However, the precise role of the FAT and whether its function is mainly related to domain-general or language-specific processes has not been fully understood. The present study aims to clarify these aspects by examining for the first time the relationship between the microstructural properties of the FAT and the performance in the first (L1) and second (L2) language in bilinguals. The comparison between unimodal and bimodal bilinguals will further allow us to understand to what extent the FAT is involved in speech control or, more generally, in language actions independent of their modality.

### 1.1 Language control in Bimodal and Unimodal bilinguals

It is widely recognized that both languages known by a bilingual person are simultaneously active, even when only one is in use (for a review, see Kroll et al., 2015). This implies that languages can influence each other and compete for output control. Therefore, learning a second language involves learning how to control and regulate competition between the second language (L2) and the native language (L1). Control in language production might be proactive and reactive; it might occur at different levels (e.g., at the language level, at word level, at the phonological level, and at the motor-articulatory level) and through different mechanisms (e.g., monitoring, shifting, inhibition). Such complex processes are orchestrated by a cortical-subcortical network, partially overlapping with the general executive control network, which involves, among other regions, the prefrontal cortex and the SMA complex. In particular, together with the anterior cingulate cortex (ACC), the pre-SMA is typically more active in bilinguals during language switching, language selection tasks, and cross-linguistic conflict resolution (for reviews, see Calabria et al., 2018; Hayakawa & Marian, 2019; Hervais-Alderman & Babock, 2020), suggesting that these clusters of regions play an important role in the language control network, likely in relation to monitoring for the correct response. The inferior prefrontal cortex (and the IFG) is more active not only during language switching but also in simple language production tasks, in particular when it comes to producing the weaker language (L2). This suggests that this region is primarily involved in response control, likely in processes of response inhibition and/or selection (Abutalebi & Green, 2016). Consistent with functional neuroimaging evidence, studies investigating structural changes associated with bilingualism suggest that ventrolateral and dorsolateral prefrontal cortices, including IFG, are consistently affected by bilingual experience (for a review, see, e.g., Pliatsikas, 2019). Furthermore, Wang & Tao (2024) directly explored the relationship between functional and structural connectivity and showed that functional connectivity within the control network significantly predicts structural connectivity between the ACC/pre-SMA and the lateral prefrontal cortex.

Control requirements during language processing in bilinguals may also depend on the language modality. Unimodal bilinguals (UBs) are individuals who have acquired two spoken languages, whereas bimodal bilinguals (BBs) have acquired a spoken and a signed language. This difference in L2 modality could be associated with differences in the way the brain controls, represents, and handles the two languages. For instance, BBs can simultaneously utter a word and make a sign, a common experience in everyday conversations (Emmorey et al., 2008). In contrast, UBs always need to select one word in one language for production. Moreover, an overwhelming preference for code blends in BBs has been observed, both during signing and speaking. Interestingly, an asymmetry has been observed in the pattern of code blends; single signs are produced frequently during speaking, whereas single words are very rarely produced when the sign language is selected as the matrix language (Emmorey et al., 2015). In summary, given that BBs often produce elements of the two languages at the same time, it has been proposed that they do not need to control the activation of the language not in use to the same extent as UBs (Emmorey et al., 2008; 2015). Contrary to this conclusion, some studies have shown a relationship between bimodal bilingualism and executive functions. Kushalnagar et al. (2010) found that highly proficient BBs performed better than less proficient BBs in an attention-switching task. Similarly, Giezen et al. (2015) reported that BBs with higher inhibitory control abilities, measured by performance in a spatial Stroop task, were less sensitive to cross-language competition than BBs with lower inhibitory control abilities.

Few previous neuroimaging studies directly investigated the effects of sign language experience on the anatomical structure of the brain, often comparing BBs or deaf signers with hearing monolinguals, thus confounding the effect of bilingualism with sign language experience. Recently, McCullough and Emmorey (2021) attempted to isolate the plasticity effects uniquely due to sign language experience comparing deaf signers Vs. hearing controls and BBs Vs. hearing controls. Sign language use was associated with a reduction in cortical thickness in the right occipital lobe and with an expansion of the surface area of the left anterior temporal lobe and the left occipital lobe. These effects were interpreted as consequences of higher demands for visual-spatial processing related to signed languages, requiring constant joint processing of motion information from the hands producing the sign and facial expressions conveying syntactic and pragmatic information. Quartarone et al. (2022) employed Diffusion Magnetic Resonance Imaging Tractography (DTI) to compare the microstructural properties of the ventral WM tracts in UBs and BBs. The results highlighted similarities and differences between the two groups. For both UBs and BBs, the degree of bilingualism was associated with the microstructural properties of the right ILF. However, only for BBs, performance on a fluency task in L1 was associated with the microstructural properties of the right Uncinate fasciculus (UF), an anterior white matter tract that connects the most anterior part of the temporal lobe with the Inferior Frontal Gyrus (IFG). This suggests that this tract could be involved in a network that controls signed L2 during L1 production. Other studies directly investigated the effect of bimodal bilingualism on frontal control regions, partially supporting the idea that BBs may rely less on executive control resources than UBs. Olulade et al. (2016) showed that the differences observed in the control regions (bilateral frontal and right parietal) between UBs and monolinguals were absent when comparing BBs and monolinguals. On the contrary, Zou et al. (2012) and Li, L. et al. (2017) found substantial structural similarities between UBs and BBs in the same areas.

In conclusion, when producing language, bilinguals constantly need to select one language and to inhibit the one not in use, likely relying on a control network involving areas that are connected by the FAT, i.e. SMA, pre-SMA complex, and the IFG. Whether the same network/areas are involved in the control of spoken and signed language remains to be fully elucidated. In the present study, our aim is to explore the role of FAT in the spoken and signed language control network.

### 1.2 The present study

We extracted the microstructural properties of the right and left FAT by means of DTI in two groups of bilinguals. We adopted the Spherical Deconvolution approach (Tournier et al., 2004; Dell’Acqua et al., 2010, 2013) and we characterized the structure of WM fibers using the Hindrance Modulated Orientational Anisotropy (HMOA), a true tract-specific index that better reflects the microstructural organization of tracts in comparison to the more classical Fractional Anisotropy (FA) measure (Dell’Acqua et al., 2013). Higher HMOA values indicate greater fiber integrity in a given direction.

To investigate the role of the FAT in bilingual language control, we explored the association between the HMOA of the left and right FAT with behavioral performance in two language production tasks and a language comprehension task, performed in both L1 and L2 by UBs and BBs. Both groups of bilinguals had a spoken language as L1 (Italian), while their L2 was English or Italian Sign Language. This approach has the advantage of highlighting what changes in WM are associated with the experience and processing of L2 according to modality, revealing both similarities and differences between the spoken and signed L2. Furthermore, by comparing the pattern observed in L1 and in L2, we could investigate the role of the FAT in selecting L2 (either signed or spoken) and/or inhibiting L1 (spoken) and the control needs to select L1 (spoken) and/or inhibiting L2 (either signed or spoken).

The two language production tasks were verbal fluency and picture naming. Picture naming is considered to simulate word production as it occurs in more naturalistic settings. After identifying the concept depicted in the picture, the corresponding lexical representation is retrieved from memory, and its phonological structure and content are specified. At this point, articulatory processes can be planned and, finally, the word is uttered. These processes do not occur automatically. Control processes are required to select the lexical item and its segments, initiate speech, and monitor output at different levels (Levelt et al., 1999; Hartsuiker & Kolk, 2001; Nozari et al., 2011; Tourville & Guenter, 2011). Only a few recent studies investigated the relationship between picture naming and the FAT. Zhong et al. (2022) estimated the integrity of the FAT in a group of patients with left post-stroke aphasia, finding that lower integrity of the FAT was associated with better picture naming performance. Troutman & Diaz (2020) showed a positive correlation between the microstructural properties of the dorsal network (which comprises the left FAT, the arcuate fasciculus, and the superior longitudinal fasciculus) and the accuracy in picture naming in the presence of phonological distractors. Higher FA and lower radial diffusivity (RD) of the dorsal pathway were associated with lower accuracy. Additionally, Troutman et al. (2022) instructed a group of healthy adults to name everyday objects and respond “picture” in the case of abstract images. They found that FA and RD of the left FAT predicted the impact of age on picture naming latency, when adjusted for associated accuracy. However, the direction of the correlations was somehow puzzling. Higher FA (but also higher RD) were linked to higher efficiency, leading to shorter adjusted latencies when naming everyday objects. In contrast, the adjusted latency for abstract images displayed an inverse pattern where higher FA and lower RD were associated with longer latencies. In conclusion, evidence suggests that the microstructural properties of the left FAT could be correlated with naming performance; however, the mixed pattern of results does not allow us to determine which are the exact processes of this task in which this tract is involved.

Verbal fluency tasks require participants to recall as many words as possible in one minute, according to specific criteria. In the semantic fluency task, participants are asked to retrieve words (or signs) within a given semantic category. The phonological fluency task requires participants to retrieve words beginning with a given phoneme or signs made with a given handshape (or other phonological parameters). Previous findings on the involvement of the FAT in verbal fluency are inconsistent: some studies reported correlations with both phonological and semantic fluency (Blecher et al. 2019; Li, M. et al. 2017), others solely with phonological fluency (Keser et al., 2020), and some found no correlation (Vallesi & Babcock, 2020). These contrasting results may be due to the multifaceted nature of the fluency task. The fluency task engages control processes across multiple levels, including actively searching among lexical items based on a single cue, selecting the item to be uttered among several activated lexical entries, sequencing the activated relevant lexical entries, and monitoring the output to prevent repetitions and invalid responses. Performance in this task hinges not only on vocabulary knowledge and speed of lexical activation, but also significantly on executive control. Luo et al. (2010; see also Sandoval et al., 2010) proposed analyzing the time course of word retrieval during the fluency task to separate the executive control component from purely lexical processes and vocabulary knowledge. Generally, participants produce many words early in the 1-minute interval, with the production rate gradually declining until reaching an asymptote. Plotting the number of items generated against time allows us to define the function representing the rate of recall. Although the exponential function has been used to describe the decline in free recall tasks (Wixted & Rohrer, 1994), research on verbal fluency has used the logarithmic function (Luo et al., 2010). The slope of the function reflects how linguistic resources are handled during the interval. With time, increased control is needed to search within the active representations in the lexicon, monitor the production of new items, resist lexical interference, and contrasting the tendency to repeat previously produced responses. Therefore, as suggested by Luo et al. (2010) and Friesen et al. (2015), the slope of the function can be viewed as an index of the control requirements during word production: the flatter the function (i.e., a slope closer to 0), the more robust the mechanism for controlling and monitoring lexical production. In our study, we used the slope of the recall rate function as an index of executive control ability in the fluency task.

The language comprehension task involved judging the acceptability of spoken or signed sentences. Participants were presented with one sentence at a time and were required to evaluate the grammatical acceptability of the sentence (acceptable / not acceptable) by pressing one of two keys. Half of the sentences were completely acceptable, while the other half contained a violation at the lexical, semantic, or syntactic level. This task served as a control task to investigate whether the neural circuit comprising the FAT exclusively handles speech/action control or whether it is also engaged in other language processes not directly associated with speech motor planning. This issue is still underexplored, and empirical evidence on this matter is still limited.

If the FAT plays a role in inhibitory language control, we might anticipate an association between the microstructural properties of this tract and the production (but not the comprehension) tasks. This correlation should manifest itself particularly during L2 production, given the stronger interference from L1 to L2 compared to the reverse. Furthermore, we expect maximum control needs for less proficient bilinguals who need to overcome the activation of their dominant L1. As the automaticity of lexical access increases for L2 and the linguistic system adapts to managing two languages, the necessity for inhibitory control process becomes less essential, and both languages can be handled with a larger degree of automaticity (Abutalebi & Green, 2007; Grundy et al., 2017; Pliatsikas, et al., 2020). Moreover, if controlling languages within the same modality engages the same neural circuit as controlling languages of different modalities, we might observe similar patterns for UBs and BBs. If instead the control of signed and spoken language relies on distinct neural circuits, UBs and BBs would exhibit different patterns, especially during L1 production when either spoken or signed language needs to be controlled.

## 2. METHOD

### 2.1 Participants

Forty-nine bilingual individuals participated in this study, which was part of a larger investigation aimed at exploring structural differences in bilingualism based on modality (refer also to Anonymous, 2022). All participants were all right-handed, confirmed through the Edinburgh Handedness Inventory Test (Oldfield, 1971) and had no history of neurological illness. Twenty-four participants had Italian as L1 and Italian Sign Language (LIS) as L2, while 25 had Italian as L1 and English as L2. UBs had a certified level of English proficiency equivalent to at least the C1 of the Common European Framework of Reference for Languages (CEFR). Additionally, within the last 5 years, they had spent at least 6 months in an English-speaking country. BBs had achieved at least the third grade of LIS proficiency, corresponding to a comprehensive mastery of the language, comparable to the C1 level of English. Both samples included 3 native bilinguals exposed to their respective L2 before the age of 3. However, the majority of the participants were sequential bilinguals who actively acquired their L2. At the time of the test, all participants reported using L2 daily.

Table 1 reports the detailed characteristics of the participants in both groups. The two samples were largely matched in all variables, except for the age of initial exposure to L2 (L2 Age of Acquisition, AoA). This is due to the fact that English is a compulsory subject in Italian primary school over the past 20 years, leading the majority of UBs to be exposed to English at around 6 or 7 years of age. On the contrary, learning LIS for people not in deaf families typically begins during adolescence, often driven by personal interests.

**Table 1.** Means and standard deviations (SD, in parentheses) of the characteristics of the two groups of bilinguals. AoA refers to the age of acquisition. The percentage of switches refers to the percentage of people who reported a given frequency of switching (from both L1 to L2 and L2 to L1).

|  | Unimodal Bilinguals, M (SD) |  |  |  |  | Bimodal Bilinguals, M (SD) |  |  |  |  |
| --- | --- | --- | --- | --- | --- | --- | --- | --- | --- | --- |
| N° | 25 |  |  |  |  | 24 |  |  |  |  |
| Gender | 8 M – 17 F |  |  |  |  | 1 M – 23 F |  |  |  |  |
| Age in years | 25.4 (4.93) |  |  |  |  | 27.79 (6.01) |  |  |  |  |
| Raven SPM | 41.61 (2.68) |  |  |  |  | 40.12 (5.39) |  |  |  |  |
| L2 AoA | 6.04 (1.54) |  |  |  |  | 16.7 (7.84) |  |  |  |  |
| Years of L2 knowledge | 18.56 (5.20) |  |  |  |  | 11.08 (10.14) |  |  |  |  |
| Self-reported proficiency | 7.2 (1.22) |  |  |  |  | 7.8 (1.88) |  |  |  |  |
| % L2 use | 47.92 (20.79) |  |  |  |  | 42.08 (24.88) |  |  |  |  |
| % of language switches | Never | Rarely | Some times | Often | Almost always/always | Never | Rarely | Some times | Often | Almost always/Always |
|  | 0 | 12 | 20 | 56 | 12 | 0 | 21 | 25 | 42 | 12 |

Participants participated in two experimental sessions conducted over two days: one session involved brain MRI and the other aimed to collect demographic and behavioral measures. The behavioral session occurred approximately a month after the scanning session. Two participants (one UB and one BB) experienced personal inconveniences, which caused the second session to be postponed by approximately 5 months. The participants received a monetary compensation of 40 euros.

### 2.2. MRI data acquisition

Diffusion imaging data was acquired using a Siemens Avanto 1.5T scanner housed in the Padova University Hospital with actively shielded magnetic field gradients (maximum amplitude 45mT/m^-1^). The body coil was used for RF transmission and an 8-channel head coil for signal reception. Protocol consisted of a localizer scan, followed by a single-shot, spin-echo, EPI sequence with the following parameters: TR = 8500, TE = 97, FOV = 307.2 x 307.2, matrix size = 128 x 128, 60 slices (no gaps) with isotropic (2.4 x 2.4 x 2.4 mm^3^) voxels. The maximum diffusion weighting was 2000 sec/mm2, and at each slice location 7 images were acquired with no diffusion gradients applied (b = 0 s/mm^2^), together with 64 diffusion-weighted images in which gradient directions were uniformly distributed in space and repeated three times, to increase signal to noise ratio. Gains and scaling factors were kept constant between acquisitions. Scanning lasted approximately 30 minutes.

#### 2.2.1. Correction of motion and eddy current distortion, and estimation of the fiber orientation distribution

Row image data from each subject were examined before proceeding on to further analyses to detect any outliers in the data, including signal drop-outs, poor signal-to-noise ratio, and image artifacts such as ghosts. Any subject whose raw data contained volumes with significant image quality issues was removed from further analysis.

DWI datasets were concatenated and corrected for subject motion and geometrical distortions using ExploreDTI (http://www.exploredti.com; Leemans et al., 2009). Spherical deconvolution (Dell’Acqua et al., 2007) approach was chosen to estimate multiple orientations in voxels containing different populations of crossing fibers (Alexander, 2007). Spherical deconvolution was calculated applying the damped version of the Richardson-Lucy algorithm with a fiber response parameter α =1.5, 400 algorithm iterations and η=0.15 and ν=15 as threshold and geometrical regularization parameters (Dell’Acqua et al., 2010). Fiber orientation estimates were obtained by selecting the orientation corresponding to the peaks (local maxima) of the FOD profiles. To exclude spurious local maxima, we applied both an absolute and a relative threshold on the FOD amplitude (Dell’Acqua et al., 2013). The first “absolute” threshold corresponding to a HMOA threshold of 0.2 was used to exclude intrinsically small local maxima due to noise or partial volume effects with isotropic tissue. This threshold was set to select only the major fiber orientation components and exclude low amplitude spurious FOD components obtained from gray matter and cerebro-spinal fluid isotropic voxels. The second “relative” threshold of 5% of the maximum amplitude of the FOD was applied to remove remaining unreliable local maxima with values greater than the absolute threshold but still significantly smaller than the main fiber orientation (Dell’Acqua et al., 2013).

#### 2.2.2 Tractography Algorithm

Whole brain tractography was performed selecting every brain voxel with at least one fiber orientation as a seed voxel. From these voxels, and for each fiber orientation, streamlines were propagated using a modified Euler integration with a step size of 0.5 mm. When entering a region with crossing white matter bundles, the algorithm followed the orientation vector of the least curvature. Streamlines were halted when a voxel without fiber orientation was reached or when the curvature between two steps exceeded a threshold of 45°. Spherical deconvolution and tractography processing were performed using StarTrack, a freely available Matlab software toolbox developed by Flavio Dell’Acqua (NatBrainLab, King’s College London), based on the methods described by Dell’Acqua et al. (2013).

#### 2.2.3 Tractography dissections of the frontal aslant tract

To visualize the frontal aslant tract and quantify tract-specific measures, we used TrackVis software (http://www.trackvis.org; Wang et al., 2007). We used the two regions of interest (ROIs) approach, according to a dissection method previously described in Budisavljević et al. (2017), Catani et al. (2012), and Rojkova et al. (2016). Two separate frontal ‘AND’ ROIs were manually delineated on the FA maps of each subject in each hemisphere. The ‘AND’ ROI is used to represent an obligatory passage for the tract, and includes the desired streamlines passing through it. We delineated on axial slices an ‘AND’ ROI around the white matter of the superior frontal gyrus (SFg ROI) and a sagittal ‘AND’ ROI around the white matter of the inferior frontal gyrus (also including the pars opercularis, triangularis and orbitalis) (IFg ROI). An example of tractography reconstructions in a representative subject is shown in Figure 1.

#### 2.3 Behavioral testing

During this session, participants completed three tasks: fluency, picture naming, and a grammaticality judgment task. Each task was performed first in L1 and then in L2. Task order remained consistent for all participants, to focus on comparing diffusion tractography measures and behavioral measures collected under identical conditions, rather than comparing performance between tasks or groups. In the semantic fluency task, participants generated words or signs within the categories “Animals” and “Transports” for Italian and “Food” and “Clothes” for English and LIS. In the phonological fluency task, the participants produced words beginning with the phonemes “F” and “L” in Italian, the phonemes “S” and “P” in English, and with the hand configurations “1” and “B” in LIS. Participants were instructed to produce words as quickly and accurately as possible in one minute while avoiding repetitions, derivatives, personal, and geographical names, which were considered errors. Responses were audio-recorded (for Italian and English) and video-recorded (for LIS). For vocal responses, participants wore Microsoft LifeChat LX-3000 earplugs equipped with a built-in microphone.

For signed responses, a camera captured the participant’s peripersonal space. The participants began the task with their hands on the table and returned to the starting position after each sign. All audio or video recordings were carefully reviewed and each word/sign was manually transcribed. To estimate the slopes of the recall rate function, the 1-minute interval was divided into 12 bins of 5 seconds each. The analysis was conducted on a participant basis. The mean of correct responses was computed for each of the 12 bins, separately for the phonological and semantic tasks in L1 and in L2. A logarithmic function was fitted to the 12 bin means and the y value of the function represented the slope value used for the analyses.

For the picture naming task, 100 colored pictures of concrete objects were selected from existing databases (Alario & Ferrand, 1999; Dell’Acqua, Lotto, & Job, 2000; Bonin, et al., 2003). The list of pictures is available in the Open Source Foundation (OSF) repository at the following link: https://osf.io/bv8fm/?view_only=beb76a1309484a64bd03727c1004f90c. The images were displayed on the computer monitor (PC Acer Intel Core i7, display 17”) one by one, centered within a white 400x400 mm template, for 2000 ms or until the participant responded. Each image was preceded by a fixation point (+) that lasted 500 ms. For presenting the pictures, we used DMDX software (Forster & Forster, 2003) for Italian and English and E-prime 2.0 (Schneider, Eschman, & Zuccolotto, 2002) for LIS. A set of 50 pictures was presented twice in each language condition in separate blocks, with a different random order for each block and participant. The blocks were separated by a short pause. The experimental blocks were preceded by a 6-trial training session. Vocal responses were recorded through the microphone. Mean RTs were manually calculated by checking the interval between the appearance of the target picture and the onset of each correct response, using Check-Vocal software (Protopapas, 2007). Manual responses were video recorder and verified for accuracy. To record the response times, at the beginning of each trial, participants were instructed to press the "Z" and "M" keys on the keyboard with their left and right index fingers. Release times of correct responses, measured from the presentation of the picture, were computed. Responses faster than 200 ms or slower than 4000 ms were considered outliers and were excluded from the RTs analysis for both vocal and manual responses. To compare manual and vocal responses, picture naming latencies in L2 were z-transformed.

For the acceptability judgment task, we created 90 correct and 90 incorrect sentences for each language, ranging from 4 to 9 words or signs each. All sentences followed the same structure, comprising a noun and a verb phrase. The stimuli are available in the OSF repository (https://osf.io/bv8fm/?view_only=beb76a1309484a64bd03727c1004f90c). The incorrect sentences concluded with either a pseudo-word/a pseudo-sign (lexical violation), or a semantically incongruent word/sign (semantic violation), or a syntactic violation. In English and Italian the syntactic violation involved a morphological violation related to number (or gender for Italian) and in LIS it involved a) a violation of the object-verb/action location concordance, or b) a violation of the negation position. For Italian and English, stimuli consisted of audio recordings of a native Italian-English bilingual, while for LIS stimuli consisted of video recordings of a native deaf signer. We presented sentences using DMDX software. Each sentence was presented after a fixation cross (“+”) lasting 1500 ms. Participants were instructed to press the “B” key for acceptable sentences and the “N” key for unacceptable ones. Since the sentences varied in durations, we calculated response latency relative to the sentence duration: RT of each correct response / sentence duration. In addition, given the different durations of oral and signed sentences, the latencies in L2 were z-transformed. Only correct responses to acceptable sentences were considered.

After the experimental session, we collected demographic data and information on the use and proficiency of L2. Following this, non-verbal intelligence was assessed by administering the Raven Standard Progressive Matrices. The research protocol received approval from the Ethics Committee for Psychological Research of the University of Padova. (Protocol n. 2015).

### 2.4 Statistical analyses

All analyses were performed with R software (R Core Team, 2020). We were interested in investigating the association between the linguistic tracts and the microstructural properties of the FAT, also in relation to the type of bilingualism (unimodal Vs. bimodal). We used regression models with the HMOA of the right/left FAT as dependent variable and group, task performance, and their interaction as predictors (Syntax: lm(FAT_L/R_HMOA ∼ Group * TaskPerformance). The effect of task performance would indicate whether the tract is involved in the processes required to accomplish the task; the interaction would highlight differences in the involvement according to the language modality. Helmert contrasts were performed, and unimodal bilinguals were set as a reference level for the models. Separate models were run for each task. To avoid false positives by multiple tests, we corrected the alpha level for statistical significance according to Bonferroni, separately for each tract and each language (alpha = 0.0125). Preliminarily, we controlled for the effects of Age of Acquisition, Gender and Age on the HMOA values. To test whether the two groups of bilingual participants showed anatomical differences in the microstructural properties of the FAT, we also added Group as predictor (Syntax: lm(FAT_L/R_HMOA ∼ Sex + Age + Group + AoA_L2).

## 3. RESULTS

Tract dissection in a representative participant is shown in Figure 1. Figure 2 reports the HMOA values separately for each tract and for each group of bilinguals.

**Figure 2.**
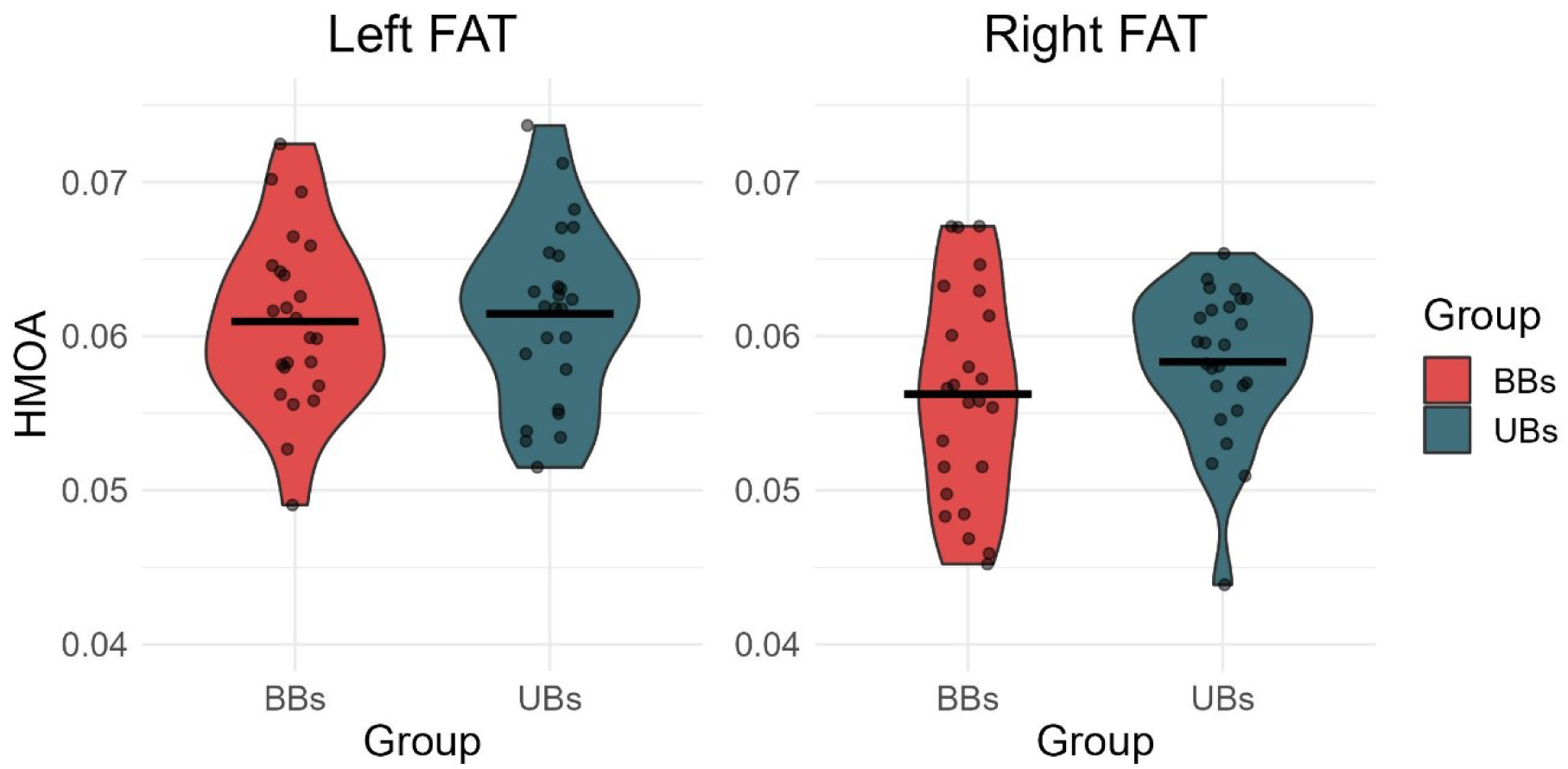
Violin plots of the HMOA values of the left and right FAT extracted from the two groups of bilinguals. Lines represent mean values.

The linear regression model run on the HMOA values did not show an effect of group (left FAT t = -0.485; right FAT t = -0.307), no effect of age (left FAT t = -0.034; right FAT t = -0.928), no effect of gender (left FAT t = -0.307; right FAT t = 0.281) and no effect of L2 AoA (left FAT t = 0.297; right FAT t = -0.443). These variables were no longer considered in the rest of the analyses.

Table 3 reports the performance of the two groups of bilinguals in behavioral tasks. A UB participant was excluded from the analysis of picture naming in L2 due to a failure in audio recording. Two UB participants were excluded from the analyses in the acceptability judgment task in L2 since their mean speed was 2.5 sd lower than the mean.

**Table 3.** Mean performance and standard deviation (in parentheses) obtained in behavioral tasks performed in L1 and L2 by unimodal (UB) and bimodal (BB) bilinguals. The Slope in the fluency task refers to the slope of the retrieval rate logarithmic function. A slope close to 0 represents a performance where control is deployed all along the minute of time. For the Grammaticality judgment task, we analyzed the responses to acceptable sentences. We used the ratio between the duration of the sentence and the latency of the keypress as an index of the response time.

|  |  | *SEMANTIC FLUENCY* |  | *PHONOLOGICAL FLUENCY* |  | *PICTURE NAMING* | *ACCEPTABILITY JUDGMENT* |  |
| --- | --- | --- | --- | --- | --- | --- | --- | --- |
|  |  | N of Words | Slope | N of Words | Slope | Mean RTS | RTs/Duration | Accuracy |
| **L1** | **BB** | 17.146 (3.447) | -1.223 (0.414) | 14.229 (2.870) | -0.686 (0.315) | 789.958 (78.867) | 1.254 (0.123) | 0.947 (0.035) |
|  | **UB** | 18.16 (2.779) | -1.272 (0.370) | 14.12 (3.153) | -0.705 (0.216) | 784.8 (91.652) | 1.213 (0.11) | 0.964 (0.032) |
| **L2** | **BB** | 14.396 (3.455) | -0.607 (0.222) | 8.875 (1.941) | -0.293 (0.177) | 1156.646 (399.056) | 1.011 (0.063) | 0.736 (0.099) |
|  | **UB** | 15.1 (2.905) | -0.905 (0.255) | 13.28 (2.292) | -0.577 (0.224) | 963.788 (125.141) | 1.367 (0.134) | 0.834 (0.068) |

The full results of the models are reported in the file “Regression tables” available online at the following OSF link: https://osf.io/bv8fm/?view_only=beb76a1309484a64bd03727c1004f90c. In the following paragraphs, we report statistically significant effects (p _Bonferroni corrected_ < .00125).

For both groups of bilinguals, the left FAT seems to be primarily involved in the semantic fluency task when performed in L2. As illustrated in Figure 3, the slope of the retrieval rate function in semantic fluency was positively correlated with the HMOA value (t=3.584, p=.0008). No significant interaction with Group was obtained (t=0.359), suggesting a similar pattern for BBs and UBs. Interestingly, the picture naming latencies in L2 also show a similar trend in both groups, even if the effect did not reach statistical significance (t=2.403, p=0.0205; see panel B, Figure 3; for the whole correlation pattern, see the file “Correlations” available in the OSF repository: https://osf.io/bv8fm/?view_only=beb76a1309484a64bd03727c1004f90c.

**Figure 3.**
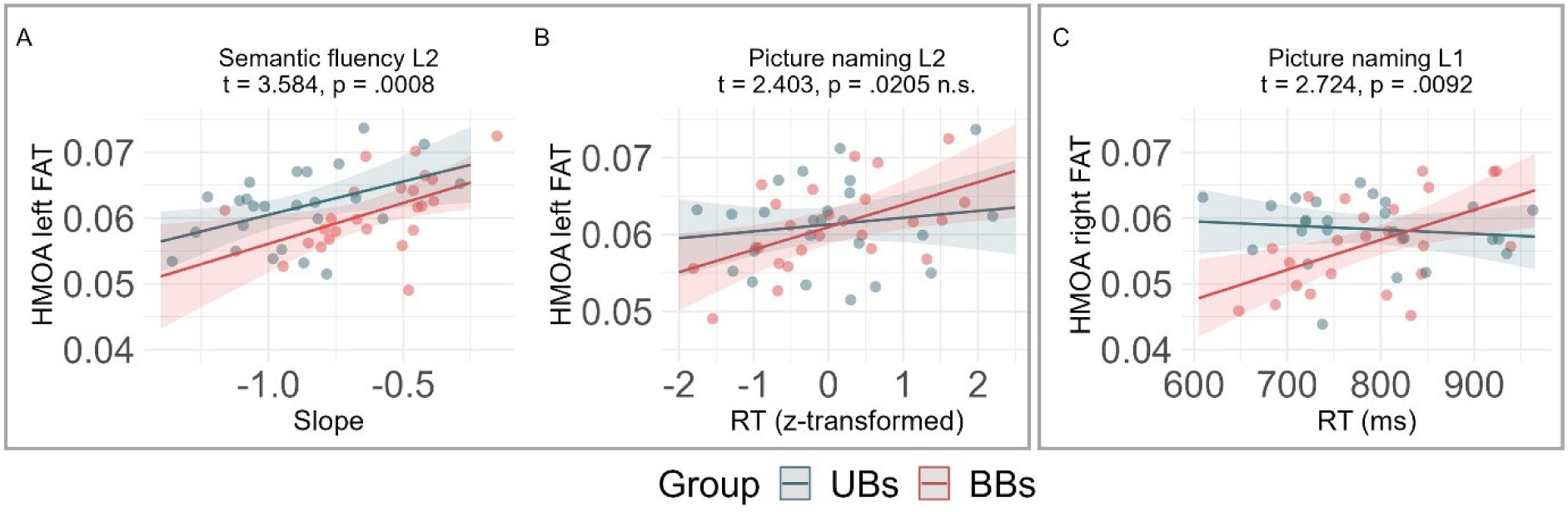
Panel A represents the model estimates for the effect of the slope of the retrieval rate function in semantic fluency in L2 on the HMOA of the left FAT. The slope increases as the HMOA increases similarly for both BBs and UBs. Panel B reports the data obtained from the analysis of picture naming latencies in L2. The effect did not reach the significance level, but there is a trend suggesting that RTs increased with increasing HMOA values. Panel C represents model estimates for the interaction between picture naming latencies in L1 and Group. For BBs only, the RTs in L1 picture naming increased as the HMOA increased.

As for the HOMA of the right FAT we obtained a significant interaction between picture naming latencies in L1 and Group (t=2.724, p=.0092). As illustrated in panel C of Figure 3, only for BBs, higher HMOA values were associated with longer picture naming latencies in L1.

Several important conclusions could be drawn from the observed pattern. First, it appears that FAT is correlated with production tasks but not with comprehension tasks. Second, production in L2 is associated with the microstructural properties of the left FAT in all participants, regardless of the modality of L2. The HMOA decreases as the slope of the recall rate function of the semantic fluency task decreases. Consistently, we found a positive trend between HMOA and picture naming latencies, indicating that lower HMOA values were associated with faster response times. Third, only for BBs, production in L1, and specifically in picture naming, is related to the properties of the right FAT. The effect is modality specific, i.e., depending on the modality of L2 during the use of L1. The neural circuit involving the right FAT appears to play a significant role during L1 production, when the activation of a signed L2 must be controlled. Note that even in this case the correlation is positive, indicating that lower HMOA values are consistently associated with faster response times.

## 4. DISCUSSION

Functional neuroimaging studies showed that multilingual speakers activate areas beyond the classical perisylvian language network. These studies suggest the involvement of a specialized control pathway that allows bilinguals to use the target language while managing interference from the unintended language (Branzi et al., 2016; Calabria et al., 2018). The key areas implicated in this process are the IFG and the SMA complex. The results of the present study underscore the involvement of the FAT, the WM tract connecting these regions, in bilingualism. By comparing unimodal and bimodal bilinguals, this study offers new insights into the role of FAT within the control network.

### 4.1 The left FAT and L2 production

Our findings reveal the involvement of left FAT in L2 production, regardless of the L2 modality. Both UBs and BBs seem to rely on the neural circuit that encompasses the left FAT during L2 word/sign production. This suggests that the type of process mediated by this tract is independent of the L2 modality. During tasks such as semantic fluency or picture naming in L2, bilinguals need to select a lexical entry in L2 while managing the interference from semantically related coactivated language entries in L1, which was spoken for both groups of bilinguals in our study. Therefore, a plausible interpretation is that the left FAT is involved in controlling/stopping inappropriate but prominent L1 (spoken). This aligns with recent studies that highlight the direct role of left FAT in speech motor control (e.g., Troutman et al., 2022; Zhong et al., 2022).

Primarily based on the involvement of the left IFG in controlled lexical retrieval and phonological activation/selection (e.g., Katzev et al., 2013; Krieger-Redwood & Jefferies, 2014; Klaus & Hartwigsen, 2019), the left FAT has been hypothesized to also play a role in these processes. The results of the present study are not consistent with this hypothesis. We observed similar left FAT involvement in L2 production for both UBs and BBs, regardless of whether the active lexical/phonological representations belong to the same (spoken) modality or different (signed Vs. spoken) modalities. Direct evidence of the involvement of FAT in lexical selection remains lacking. Zyranov et al. (2020) examined the effect of FAT volume on lexical selection in post-stroke patients using a picture-word interference task and failed to find a relationship between the two variables. It could be possible that activation of the left IFG in lexical and phonological selection is mainly related to activation in temporal areas or to subcortical structures, as suggested by recent studies investigating the pattern of functional connectivity during a covert naming task (Rivas-Fernandez et al., 2021) and a picture naming task in a language switch paradigm (Wang & Tao, 2024).

Notably, we found a positive correlation between the HMOA of the left FAT and the slope of the word retrieval rate function in the semantic fluency task. The slope reflects the rate of declining in performance during the minute allowed for word/sign retrieval. Higher HMOA values are associated with slow decline rates, that is, with superior control abilities. Furthermore, a positive relationship has been observed between the HMOA of the left FAT and the L2 naming times. Longer naming times in L2 corresponded to higher HMOA values. According to our predictions, this pattern suggests that less proficient bilinguals strongly rely on active control of the native language while speaking/signing in L2. As lexical access in L2 becomes more automatic and the linguistic system adapts to using multiple languages, inhibitory control process may become less essential and/or control may be displaced at other (earlier) levels of processing. These findings align with the views that envision structural changes related to bilingualism as the result of a learning process. Early learning stages, when processes have not yet been automatized, can entail more pronounced changes in the frontal areas of the brain involved in control and executive functions (Green & Abutalebi, 2013). However, as proficiency increases and L2 lexical activation/selection becomes more and more automatic, frontal involvement is not yet required and changes in this part of the brain progressively disappear, in favor of changes in most posterior parts of the brain (Grundy et al., 2017). Consistent with this idea, Quartarone et al. (2022) found for both UBs and BBs a correlation between the proficiency/amount of use of L2 and the microstructural properties of the Inferior Longitudinal fasciculus, a ventral WM tract connecting the occipital and the temporal lobes.

Although we found significant correlations between the microstructure of the left FAT and the performance of L2 semantic fluency, no such correlations were found for phonological fluency in L2. Typically, phonological fluency is thought to strongly reflect executive control, given that the search based on a phonological cue is more artificial than the search based on a semantic cue. However, the task might be highly demanding in L2, with control resources primarily allocated to the search for the target words/signs rather than inhibiting competing candidates and thus involving neural circuits that do not comprise the FAT.

### 4.2. The role of the FAT in L1 production

Our findings revealed the involvement of the FAT in L1 production for BBs. We observed that increased HMOA values corresponded to longer response times in picture naming for this specific group of bilinguals. This pattern suggests that BBs exert specific control over L2 (signed) when naming pictures in L1. Differently from UBs, who need to select one phonological (spoken) representation at a time, BBs may activate both word and sign phonology upon the presentation of a picture, without experiencing between-language competition until later processing stages. However, a point arises when the spoken modality must be selected, and/or the signed modality must be inhibited. This process appears to involve a neural circuit involving the FAT. It is possible that this tract contributes to the control of hand movements, stopping the activation of motor plans related to the phonology of signs. Consistent with this hypothesis, Budisavljević et al. (2017) showed an association between the bilateral FAT microstructure and the hand kinematics during grasping and reaching movements performed with the right hand. This proposal is consistent with recent literature indicating the involvement of FAT in selecting and controlling appropriate motor actions (Dick et al., 2019; Shekari & Nozari, 2022). Similarly to the findings in L2, we observed positive correlations between the L1 naming times and the HMOA values. This suggests that highly proficient bilinguals maintain their language output without – or with minimal - interference from the language not in use.

Quartarone et al. (2022) reported a significant correlation between L1 production and the microstructural properties of the right UF. Parallel to the results reported here, a significant relation was present for BBs, but not for UBs. This convergence might point to some neural adaptation of the (right) frontal regions/networks to control the signed modality in BBs, consistent with evidence showing differences in control needs between BBs and UBs (for a review, see Emmorey et al., 2015).

The fact that for BBs, inhibition of the signed modality involves the right FAT, while inhibition of the spoken modality primarily engages the left FAT, further supports the idea that, on the left side, the FAT specializes in the control of speech actions (Dick et al., 2019, Shekari & Nozari, 2023; Ribeiro et al., 2024).

Previous evidence supporting the involvement of FAT in language production comes primarily from studies involving clinical populations (e.g., Basilakos et al. 2014; Blecher et al., 2019; Catani et al. 2013; Keser et al., 2020; Kinoshita et al. 2015; Li, M. et al., 2017). Conversely, research involving healthy participants failed to establish a direct association between native language word production and the microstructure of FAT (Babock & Vallesi, 2020; Kronfeld-Duenias et al., 2016). Similarly, in our study, we did not find correlations between the microstructural properties of the FAT and the performance in the L1 production tasks for UBs. For UBs (or monolinguals), when speaking in their native language, lexical selection among competing units might occur before the activation of motor plans in most cases. Inhibitory control may be applied during the activation of lexical/phonological representations through neural circuits not involving the FAT (for a review, see de Zubicaray & Piai, 2019). Alternatively, it could be proposed that when a language is fully acquired and proficiently handled, lexical selection may not necessitate inhibitory control mechanisms and the most active phonological unit is readily selected for articulation (Dell, 1986; Costa, 2005).

### 4.3 Concluding remarks

For the first time, the current study presents a direct link between the microstructure of bilateral FAT and language control during language production in bilingual individuals. No evidence of the involvement of the FAT in the acceptability judgment comprehension task was found, implying a predominant role of this tract in processes associated with bilingual language production. By comparing unimodal and bimodal bilinguals, we could further delineate the nature of the control mechanism supported by the FAT.

Our findings support the hypothesis that the FAT is mainly involved in action control. Specifically, this tract might have a role in resolving the competition among motor programs of language production, likely by inhibiting the non-target action (Dick et al., 2019; Shekari & Nozari, 202). Production performance of a spoken or a signed L2 is correlated with the microstructural properties of the left FAT. In addition, the correlation observed between the microstructure of FAT and production in L1 in BBs – but not in UBs – suggests that when languages activate different modalities, control over the unintended language might occur at later processing levels than when competition occurs between representations sharing the same phonological format. Our study also revealed some hemispheric specialization in bilingual language control. The left FAT appears to primarily manage activation of the dominant spoken language during L2 word or sign production, while the right FAT appears to be involved in controlling the signed language during speech. Furthermore, language competition appears stronger for less proficient bilinguals, suggesting that in more balanced bilinguals, the control network adapts so that competition is either mitigated or resolved before activating motor representations, likely involving other brain regions and/or other tracts of the neural network engaged in language control.

The present study has some limitations, mainly due to the challenges in pairing BBs and UBs based on the year of first exposure to L2. Typically, hearing individuals not born to deaf parents encounter sign language during adolescence, while learning a spoken L2 often begins earlier, during primary school. Consequently, the two samples did not match fully for this variable. Despite this, we believe it unlikely that AoA confounded the reported effects, as this variable did not show an impact on the microstructural properties of the FAT.

## DATA AVAILABILITY

The materials used for the behavioral tasks and the data with the identifying information removed are available at https://osf.io/bv8fm/?view_only=beb76a1309484a64bd03727c1004f90c

To request the full data set contact

## Acknowledgements

The authors thank Elena Pretato for her help with the scoring of Sign Language responses.

## Author contribution statement

CQ, FP, EN, SB contributed to the conceptualization of the study. CQ and EN developed the materials for the behavioral tasks, CQ collected the data, SB performed the DTI analysis, SG performed statistical analyses and data visualization, FP coordinated the teamwork and drafted the manuscript. All authors provided critical feedback and read and approved the final manuscript.

